# High levels of truncated RHAMM cooperate with dysfunctional p53 to accelerate the progression of pancreatic cancer

**DOI:** 10.1101/2021.02.19.432042

**Authors:** Jennifer Feng, Anthony Lin, Xiang Chen, Dunrui Wang, Megan Wong, George Zhang, Joseph Na, Tiantian Zhang, Zhengming Chen, Yao-Tseng Chen, Yi-Chieh Nancy Du

**Author notes:** Corresponding author: Yi-Chieh Nancy Du, Department of Pathology and Laboratory Medicine, Weill Cornell Medicine, New York, NY 10065. equal contribution (Jennifer Feng, Anthony Lin).

## Abstract

Pancreatic cancer has the lowest survival rate in all types of cancer. Pancreatic cancer patients are often diagnosed at advanced stages. A better therapeutic development for this devastating disease is urgently needed. Receptor for hyaluronan-mediated motility (RHAMM), not expressed in adult pancreas, has been suggested as a prognostic factor and a potential therapeutic target for pancreatic ductal adenocarcinoma (PDAC) and pancreatic neuroendocrine tumor (PNET). In this study, we initially sought to determine whether genetic deletion of *RHAMM* would slow down pancreatic cancer progression using *Rhamm*^*-/-*^ mice. However, we found that *Rhamm*^*-/-*^ mice expressed a truncated HMMR^Δexon8-16^ protein at higher abundance levels than wild-type RHAMM. While HMMR^Δexon8-16^ did not enable malignant progression of pancreatic intraepithelial neoplasia in *p48-Cre*; *LSL-KRAS*^*G12D*^ mice, it accelerated the formation of invasive PDAC and shortened the survival of *p48-Cre*; *LSL-KRAS*^*G12D*^ mice with heterozygous p53 knockout. *Kras*^*G12D*^ PDAC mice with homozygous p53 knockout mice died around 10 weeks, and the effect of HMMR^Δexon8-16^ was not apparent in these mice with short-life span. In addition, HMMR^Δexon8-16^ shortened the survival of PNET-bearing *RIP-Tag* mice, which had inactivated p53. In our analysis of TCGA dataset, pancreatic cancer patients with mutant *TP53* or loss of one copy of *TP53* had higher *RHAMM* expression, which, combined, predicted worse outcomes. Taken together, by collaborating with dysfunctional p53, high levels of HMMR^Δexon8- 16^ that lacks the centrosome targeting domain and degrons for interaction with the Anaphase-Promoting Complex (APC) accelerated pancreatic cancer progression.

## Introduction

Pancreatic cancer is the leading cause of cancer-related death. In contrast to the steady increase in survival observed for most cancer types, advances have been slow for pancreatic cancer, which is typically diagnosed at an advanced stage [1]. About 90% of all malignant pancreatic tissues are pancreatic ductal adenocarcinoma (PDAC) with a five-year survival rate of <10% [2]. *KRAS, CDKN2A, TP53*, and *SMAD4* are the four frequently mutated genes that characterize PDAC [3]. Activating *KRAS* mutations are present in more than 99% of pancreatic intraepithelial neoplasia (PanIN) and over 90% of PDAC, and has been shown to be an initiating event of PDAC using genetically engineered mouse models [4]. Studies from mouse models of PDAC further demonstrated that loss of functional p53 cooperates with mutant KRAS to cause rapid PDAC progression and metastasis [5].

Pancreatic neuroendocrine tumor (PNET) is the second most common malignancy of the pancreas. The incidence of PNET is increasing and PNET presents with a wide variety of clinical manifestations and unfavorable survival rates [6]. Unlike PDAC, the major molecular driver for PNETs is not well-understood. Recent molecular characterization of PNETs reveals alterations in pathways/genes such as mTOR pathway, *cyclin D1*/*Cdk4*/*retinoblastoma* (*Rb*), *TP53, MEN1, DAXX, ATRX, UCHL1, RHAMM*, and microRNAs [7–9]. In the *RIP-Tag* mouse model of PNET, the rat insulin promoter (RIP) drives the expression of SV40 T antigen (Tag), providing the driving force for tumor initiation by inhibiting the activities of tumor suppressors, p53 and Rb [10]. Preclinical trials in the *RIP-Tag* mice have predicted that sunitinib and everolimus would be effective in treating human PNETs [11–13]. However, sunitinib and everolimus only extend the median patient survival by roughly 6 months, and all patients eventually develop resistance to both drugs [14, 15].

Receptor for hyaluronic acid-mediated motility (RHAMM or HMMR) was identified as a receptor of hyaluronan [16], and its expression peaks at G2/M [17, 18]. Studies have shown that RHAMM plays an important role in cell motility [19]. Human *RHAMM* encodes 18 exons and alternative splicing yields four isoforms [20]. Of the four isoforms (*RHAMMv1-4*), we have shown mRNA expression levels of *RHAMMv3*, also known as *RHAMM*^*B*^, to be the most prominent in human PDAC, PNET, and liver metastases of PNET [21]. Using the *RIP-Tag; RIP-tva* mouse model of PNET and cell line xenograft mouse models, we have discovered RHAMM^B^ as the first protein that is able to promote PNET metastasis to the liver [21, 22], and we have also found *RHAMM*^*B*^ expression to correlate with poor survival in PDAC patients [21]. Most normal human tissues, including pancreas, do not express RHAMM [17, 21], and the differential RHAMM expression in normal tissues and pancreatic cancer may thus provide novel therapeutic possibilities by targeting RHAMM. To investigate whether RHAMM is a potential therapeutic target in pancreatic cancer, we set out to determine the effects of RHAMM deletion on the progression and survival of PDAC and PNET in mouse models using *Rhamm*^*-/-*^ mice [23]. However, after analyzing RHAMM mRNA and protein expression in *Rhamm-/-* mice, we unexpectedly discovered that a truncated HMMR^Δexon8-16^ protein was expressed in *Rhamm*^*-/-*^ mice and was more abundant than the full-length protein in RHAMM wide-type (WT) mice. Based on these findings, we propose to re-name *Rhamm*^*-/-*^ mice as *HMMR*^*Δexon8-16/Δexon8-16*^ mice. In this study, we investigated the effects of the HMMR^Δexon8-16^ protein on tumor progression and survival of PDAC and PNET in mouse models.

## Results

### Detection of HMMR^Δexon8-16^ protein in *Rhamm*^*-/-*^ mice

Mouse RHAMM/HMMR has 18 exons (NM_013552.2). The *Rhamm*^*-/-*^ mouse strain was previously generated by deleting exons 8 ∼16 of the *HMMR* gene through homologous recombination in embryonic stem (ES) cells [23]. It was reported that no HMMR protein was detected in the spleen lysate of *Rhamm*^*-/-*^ mice using RHAMM polyclonal antibody in Western blot analysis [23]. However, because the polyclonal antibody was generated against recombinant RHAMM protein [23] and the specific immunogenic epitope(s) are unknown, we decided to investigate whether any truncated RHAMM protein was expressed in *Rhamm*^*-/-*^ mice using a RHAMM-specific monoclonal antibody against the N-terminus (clone EPR4054). The immunogen for this clone EPR4054 was the first 50 amino acids of human RHAMM, and the antibody cross-reacted with mouse RHAMM. Full-length WT mouse *RHAMM* encodes 794 amino acids with a predicted molecular weight of ∼92 kDa. We have previously reported that the testis has the most abundant RHAMM proteins among all adult human tissues [17], so we predicted that RHAMM would also be highly expressed in mouse testis and therefore the testis is a better tissue than spleen to evaluate whether a truncated RHAMM protein was expressed. To verify that exons 8 ∼ 16 were deleted in the *Rhamm*^*-/-*^ mice, we isolated mRNA from the testis of *Rhamm*^*-/-*^ mice (n = 3), made cDNA, amplified *RHAMM* fragments by polymerase chain reaction (PCR), and performed DNA sequencing. Our analysis confirmed a fusion of exon 7 and exon 17, which resulted in a stop codon in exon 17 and would yield a truncated RHAMM of 239 amino acids with a predicted molecular weight of ∼27 kDa (Figure 1A). In addition, we found an adenine to cytosine missense mutation at nucleotide position 290, which changed the amino acid from lysine to threonine (residue 71) (Figure 1A and Figure 2). Protein lysates and histological sections from testes of WT mice and *Rhamm*^*-/-*^ mice were then evaluated for the expression of RHAMM proteins. Western blot analysis using EPR4054 antibody detected full-length RHAMM protein (∼92 kDa) in WT mouse testes and truncated RHAMM protein (∼27 kDa) in *Rhamm*^*-/-*^ mouse testes (Figure 1B). Immunohistochemical staining using EPR4054 antibody also revealed RHAMM protein in *Rhamm*^*-/-*^ mouse testes (Figure 1C). Both Western blot and immunohistochemical analyses showed that the truncated RHAMM protein (HMMR^Δexon8-16^) was highly expressed in *Rhamm*^*-/-*^ mice, at levels more abundant than WT RHAMM protein in the control WT mice. Therefore, we renamed this mouse strain from *Rhamm*^*-/-*^ to *HMMR*^*Δexon8-16/Δ exon8-16*^.

**Figure 1.**
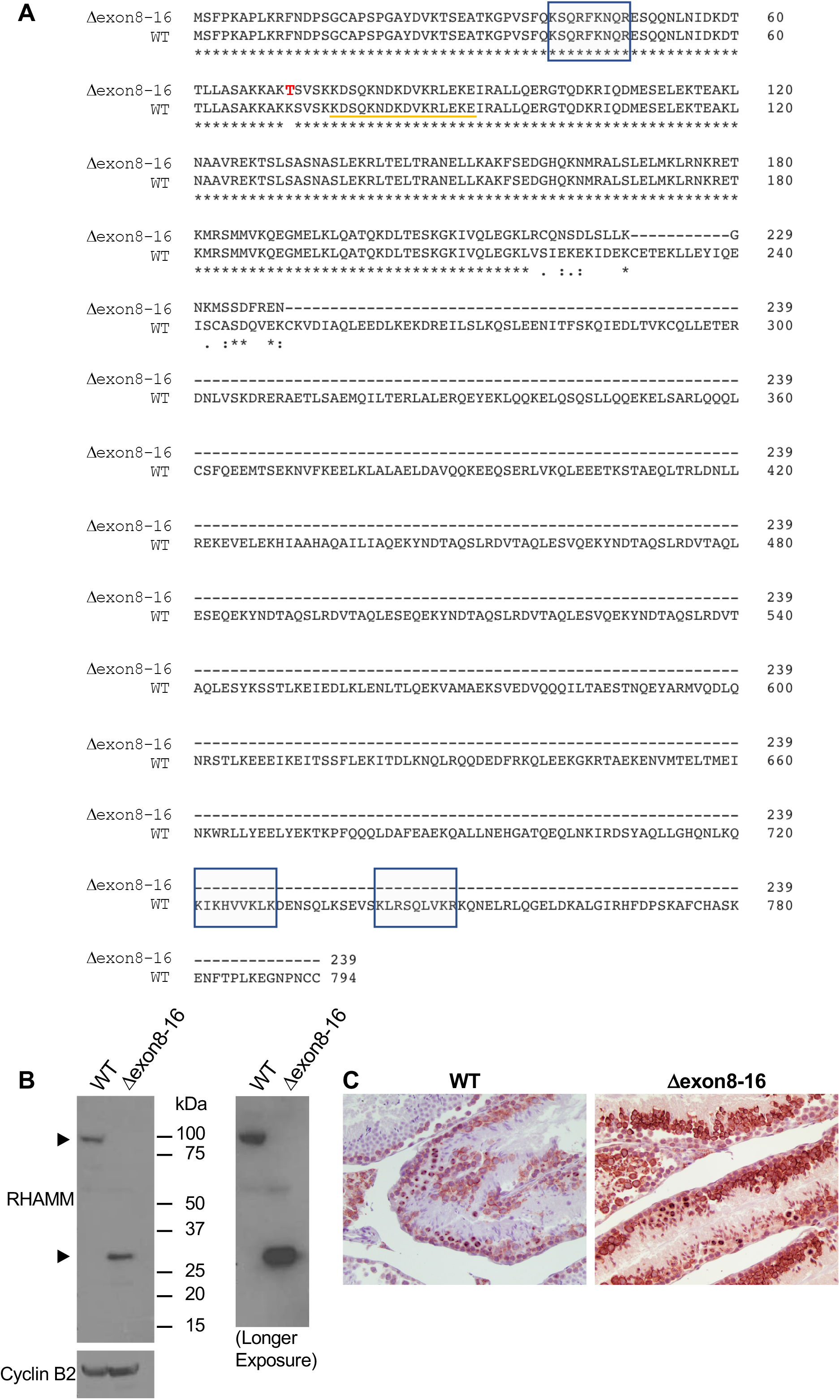
Comparison of WT mouse RHAMM/HMMR protein and HMMR^Δexon8-16^. (A) Deduced amino acid sequence of HMMR^Δexon8-16^ and WT mouse RHAMM/HMMR (translated from NM_013552.2). Potential hyaluronan-binding sites (BX7B motif) are shown in shaded boxes. Sequences from exon 4 are indicated by an orange line. A missense mutation from lysine (K) to threonine (T) at residue 71 is shown in red font. (B) Western blot analysis of mouse RHAMM in the testis of WT and *HMMR*^*Δexon8-16/Δexon8-16*^. The right panel was a longer exposure than the left panel for RHAMM. Cyclin B2 was used as a control. (C) Immunohistochemical staining showed RHAMM staining in the germ cells of testis from a *HMMR*^*Δexon8-16/Δexon8-16*^ mouse, at stronger intensity than that observed in the testis of a WT mouse. Original magnification, 200X.

**Figure 2.**
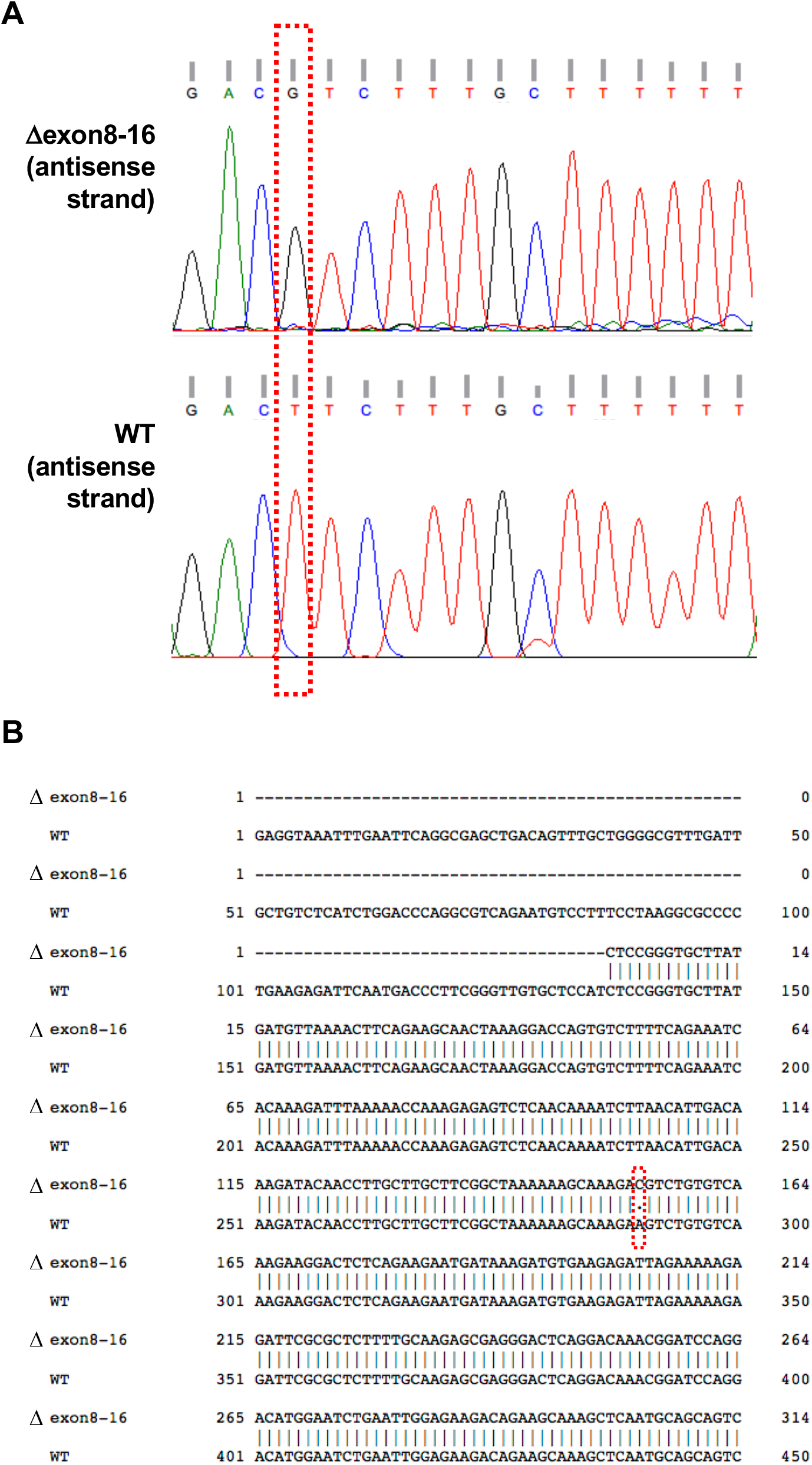
A missense mutation at nucleotide position 290 of the coding region of *HMMR*^*Δexon8-16*^. (A) DNA sequencing showed a mutation from T to G (antisense strand) from HMMR transcripts in the testis of a *HMMR*^*Δexon8-16/Δexon8-16*^ mouse, as compared to transcripts derived from the testis of a WT mouse. (B) Sequence alignment between a partial fragment of *HMMR*^*Δexon8-16*^ and a partial coding sequence of NM_013552.2 showed an A to C missense mutation at nucleotide position 290.

### *HMMR*^*Δexon8-16/Δ exon8-16*^ mice have normal pancreas development

Consistent with the previous report [23], we found no embryonic lethality for *HMMR*^*Δexon8-16/Δ exon8-16*^ mice. The appearance of *HMMR*^*Δexon8-16/Δ exon8-16*^ mice was normal throughout their lifespan and live longer than 1 year without any spontaneous tumorigenesis (data not shown). To determine whether the truncated RHAMM protein (HMMR^Δexon8-16^) causes any abnormality in pancreatic development, the pancreas from adult *HMMR*^*Δexon8-16/Δ exon8-16*^ was examined and no histologic abnormality was identified (Figure 3A).

**Figure 3.**
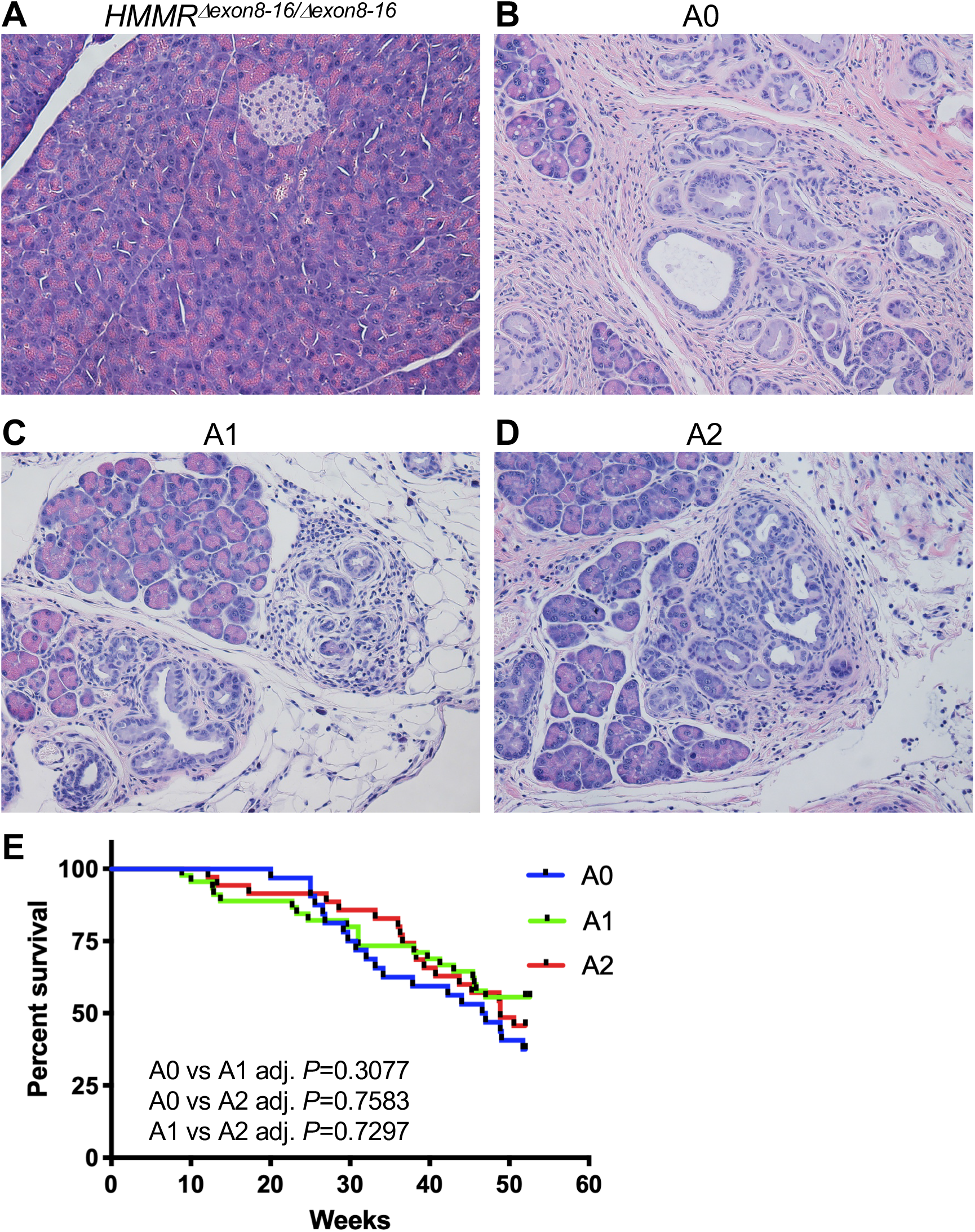
*HMMR*^*Δexon8-16/Δexon8-16*^ mice have normal pancreas and *HMMR*^*Δexon8-16*^ does not significantly change the survival of *p48-Cre*; *LSL-KRAS*^*G12D*^ mice. (A-D) Hematoxylin and eosin (H&E) stain of representative pancreas from (A) *HMMR*^*Δexon8-16/Δexon8-16*^ mouse, (B) A0: *p48-Cre*; *LSL-KRAS*^*G12D*^ mouse, (C) A1: *p48-Cre*; *LSL-KRAS*^*G12D*^; *HMMR*^*Δexon8-16/WT*^ mouse, (D) A2: *p48-Cre*; *LSL-KRAS*^*G12D*^; *HMMR*^*Δexon8-16/Δ exon8-16*^ mouse. A0, A1 and A2 mice all showed localized ductal proliferation consistent with PanINs, but no evidence of invasive PDAC. (E) Kaplan-Meier survival curve for A0, A1, and A2 mice. Tukey adjusted *P* values from pairwise log-rank test were shown. Original magnification, 200X.

### *HMMR*^*Δexon8-16*^ did not alter the course of PanIN formation in *p48-Cre*; *LSL-KRAS*^*G12D*^ mice

It has been shown that the expression of mutant *KRAS*^*G12D*^ from the endogenous locus in the mouse pancreas leads to the development of premalignant ductal lesions, PanINs, in *Pdx1-Cre*; *LSL-KRAS*^*G12D*^ mice and *p48-Cre*; *LSL-KRAS*^*G12D*^ mice [4]. Subsequent studies revealed that p53 constrains the progression of PanIN to advanced PDAC [5, 24]. *Pdx1-Cre*; *LSL-KRAS*^*G12D*^ mice harboring heterozygous p53 loss or homozygous-null p53 developed PDAC with an average latency of 21.8 weeks or 6.2 weeks, respectively [5]. Because the expression of the Cre recombinase under p48 promoter is restricted to mouse pancreas compared to that under the Pdx1 promoter [3], we decided to use *p48-Cre* instead of *Pdx1-Cre* to target mutant Kras^G12D^ expression in the pancreas in this study. To determine the effect of N-terminal RHAMM (HMMMR^Δexon8-16^) on the initiation and the progression of PanINs and pancreatic ductal adenocarcinomas (PDAC), we crossed *HMMR*^*Δexon8-16/Δ exon8-16*^ to *p48-Cre*; *LSL-KRAS*^*G12D*^ mice with WT p53 (Group A), heterozygous p53 loss (*p53*^*lox/+*^, Group B), or homozygous-null p53 (*p53*^*lox/lox*^, Group C) (Table 1). Within each group (A, B, and C), we evaluated the impact of WT HMMR (A0, B0, and C0), one copy of *HMMR*^*Δ exon8-16*^ (*HMMR*^*Δ exon8-16/WT*^: A1, B1, and C1), and two copies of *HMMR*^*Δexon8-16*^ (*HMMR*^*Δexon8-16/Δ exon8-16*^: A2, B2, and C2) (Table 1).

**Table 1.** Histological Phenotypes and Average Life Span of Mice

| Group | Genotype | Histology | Median survival (week) |
| --- | --- | --- | --- |
| A0 | *p48-Cre; LSL-KRAS<sup>G12D</sup>* | PanIN | 46.79 |
| A1 | *p48-Cre; LSL-KRAS<sup>G12D</sup>; HMMR<sup>Δexon8-16/WT</sup>* | PanIN | undefined |
| A2 | *p48-Cre; LSL-KRAS<sup>G12D</sup>; HMMR<sup>Δexon8-16/Δ exon8-16</sup>* | PanIN | 48.86 |
| B0 | *p48-Cre; LSL-KRAS<sup>G12D</sup>; p53<sup>lox/+</sup>* | PDAC | 23.93 |
| B1 | *p48-Cre; LSL-KRAS<sup>G12D</sup>; p53<sup>lox/+</sup>; HMMR<sup>Δexon8-16/WT</sup>* | PDAC | 17.50 |
| B2 | *p48-Cre; LSL-KRAS<sup>G12D</sup>; p53<sup>lox/+</sup>; HMMR<sup>Δexon8-16/Δ exon8-16</sup>* | PDAC | 15.41 |
| C0 | *p48-Cre; LSL-KRAS<sup>G12D</sup>; p53<sup>lox/lox</sup>* | PDAC | 9.14 |
| C1 | *p48-Cre; LSL-KRAS<sup>G12D</sup>; p53<sup>lox/lox</sup>; HMMR<sup>Δexon8-16/WT</sup>* | PDAC | 9.86 |
| C2 | *p48-Cre; LSL-KRAS<sup>G12D</sup>; p53<sup>lox/lox</sup>; HMMR<sup>Δexon8-16/Δ exon8-16</sup>* | PDAC | 9.57 |
| T0 | *RIP-Tag* | PNET | 17.57 |
| T1 | *RIP-Tag; HMMR<sup>Δexon8-16/WT</sup>* | PNET | 16.14 |
| T2 | *RIP-Tag; HMMR<sup>Δexon8-16/Δ exon8-16</sup>* | PNET | 15.50 |

As expected, our cohort of *p48-Cre*; *LSL-KRAS*^*G12D*^ mice (A0) developed PanINs (Figure 3B). The majority of the A group mice lived longer than 40 weeks of age (Figure 3E). We found no significant survival difference among A0, A1, and A2 after following them up to 1 year (Figure 3E, Log-rank test, A0: 32 mice, A1: 45 mice, and A2: 35 mice). Similar results were obtained from Cox regression (data not shown). Mice in A0, A1, and A2 at the end point (1 year of age) all developed PanINs, but no invasive PDAC (Figure 3B-D). Our data suggest that N-terminal RHAMM protein (HMMR^Δexon8-16^) cannot promote the progression of PanINs initiated by *KRAS*^*G12D*^ to PDAC.

### *HMMR*^*Δexon8-16*^ shortened the survival and accelerated PDAC formation in *p48-Cre*; *LSL-KRAS*^*G12D*^; *p53*^*lox/+*^ mice

We established a cohort of B group, *p48-Cre*; *LSL-KRAS*^*G12D*^ mice with heterozygous p53 loss (*p53*^*lox/+*^) and different copy numbers of *HMMR*^*Δexon8-16*^ (B0, B1, and B2). Mice in the B group lived less than a year, and all mice in these three subgroups developed advanced PDAC at death or when they were sick (Figure 4 and data not shown). Having either one copy or two copies of *HMMR*^*Δexon8-16*^ significantly shortened the survival of *p48-Cre*; *LSL-KRAS*^*G12D*^; *p53*^*lox/+*^ mice (Figure 4A, Log-rank test, B0: 86 mice, B1: 57 mice, and B2: 36 mice). The median survival ages for B0, B1, and B2 were 22.93 weeks, 17.5 weeks, and 15.14 weeks, respectively (Table 1). Cox regression analysis showed that B1 is 164% more likely to die at any time point than B0 (*P* < 0.0001) and B2 is 234% more likely to die than B0 (*P* < 0.0001). At the terminal stage of each subgroup, there was no significant different PDAC histology found between *p48-Cre*; *LSL-KRAS*^*G12D*^; *p53*^*lox/+*^ mice with either one copy or two copies of *HMMR*^*Δexon8-16*^ (Figure 4B-G, and data not shown). However, direct invasion of PDAC into peripancreatic lymph nodes and metastasis to liver were observed in mice with *HMMR*^*Δexon8-16*^ (Figure 4F and 4G).

**Figure 4.**
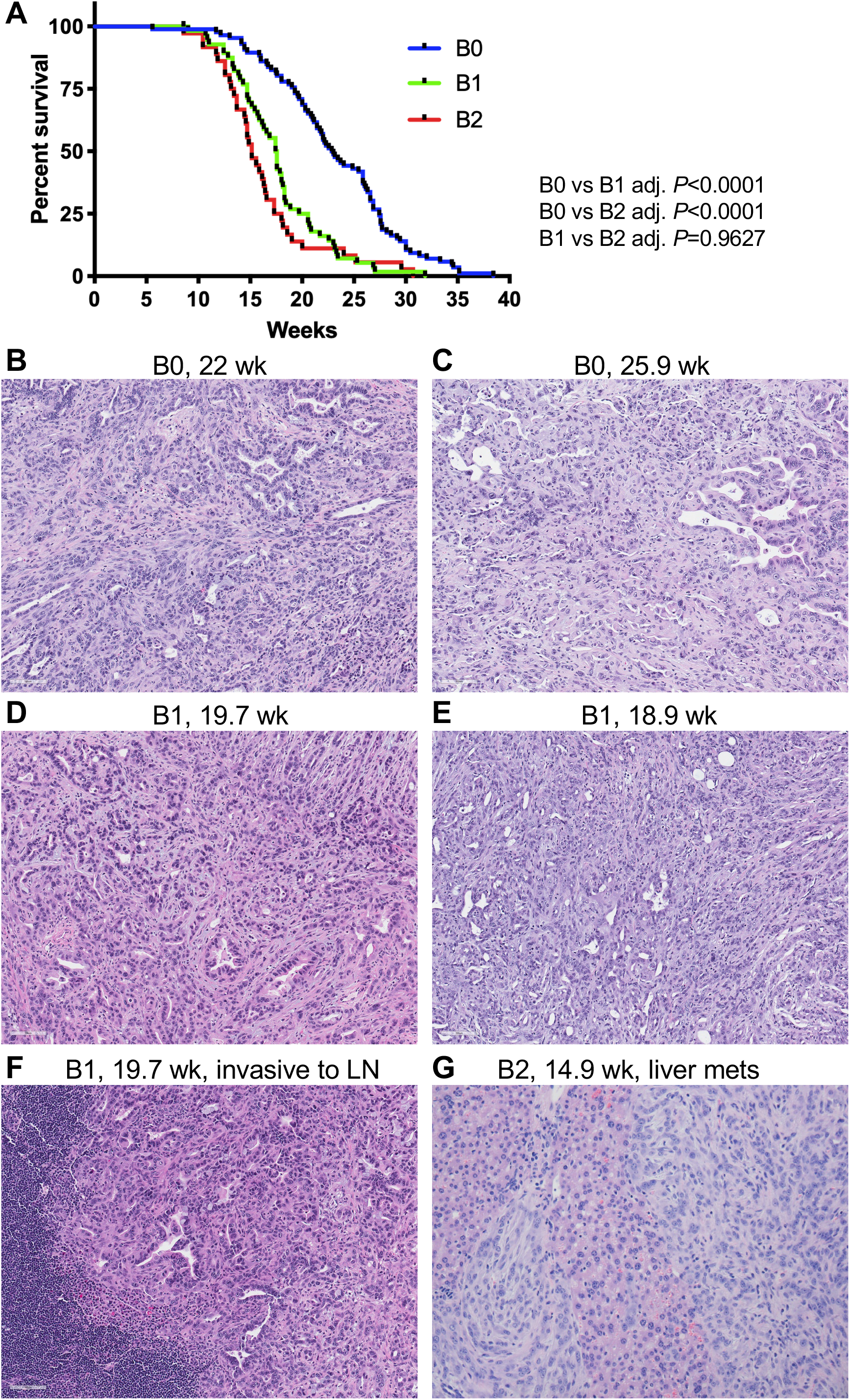
*HMMR*^*Δexon8-16*^ shortened the survival of *p48-Cre*; *LSL-KRAS*^*G12D*^; *p53*^*lox/+*^ mice. (A) Kaplan-Meier survival curve for B0, B1, and B2 mice. Tukey adjusted *P* values from pairwise log-rank test were shown. (B-E) H&E stain of invasive PDAC from (B) a 22-week-old B0: *p48-Cre*; *LSL-KRAS*^*G12D*^; *p53*^*lox/+*^ mouse, (C) a 25.9-week-old B0: *p48-Cre*; *LSL-KRAS*^*G12D*^; *p53*^*lox/+*^ mouse, (D) a 19.7-week-old B1: *p48-Cre*; *LSL-KRAS*^*G12D*^; *p53*^*lox/+*^; *HMMR*^*Δexon8-16/WT*^ mouse, and (E) a 18.9-week-old B1: *p48-Cre*; *LSL-KRAS*^*G12D*^; *p53*^*lox/+*^; *HMMR*^*Δexon8-16/WT*^ mouse. (F) H&E stain of a peripancreatic lymph node (LN) invaded by PDAC from the same animal in (D). (G) H&E stain of liver metastasis of PDAC from a 14.9-week-old B2: *p48-Cre*; *LSL-KRAS*^*G12D*^; *p53*^*lox/+*^; *HMMR*^*Δexon8-16/Δ exon8-16*^ mouse. Original magnification, 200X.

To investigate why *p48-Cre*; *LSL-KRAS*^*G12D*^; *p53*^*lox/+*^; *HMMR*^*Δexon8-16/WT*^ (B1) died earlier than *p48-Cre*; *LSL-KRAS*^*G12D*^; *p53*^*lox/+*^ (B0), we sacrificed a cohort of B0 and B1 mice at the age of 15 weeks (n =3 per group). We found that whereas B0 mice showed only PanINs and not PDAC at 15 weeks (Figure 5A and 5B), B1 mice already developed invasive PDAC (Figure 5C, 5D, and 5E). Taken together, in the presence of heterozygous p53 loss, having one copy of *HMMR*^*Δexon8- 16*^ was sufficient to accelerate progression of PanINs into invasive PDAC, leading to the shortened survival in the *p48-Cre*; *LSL-KRAS*^*G12D*^; *p53*^*lox/+*^ mice.

**Figure 5.**
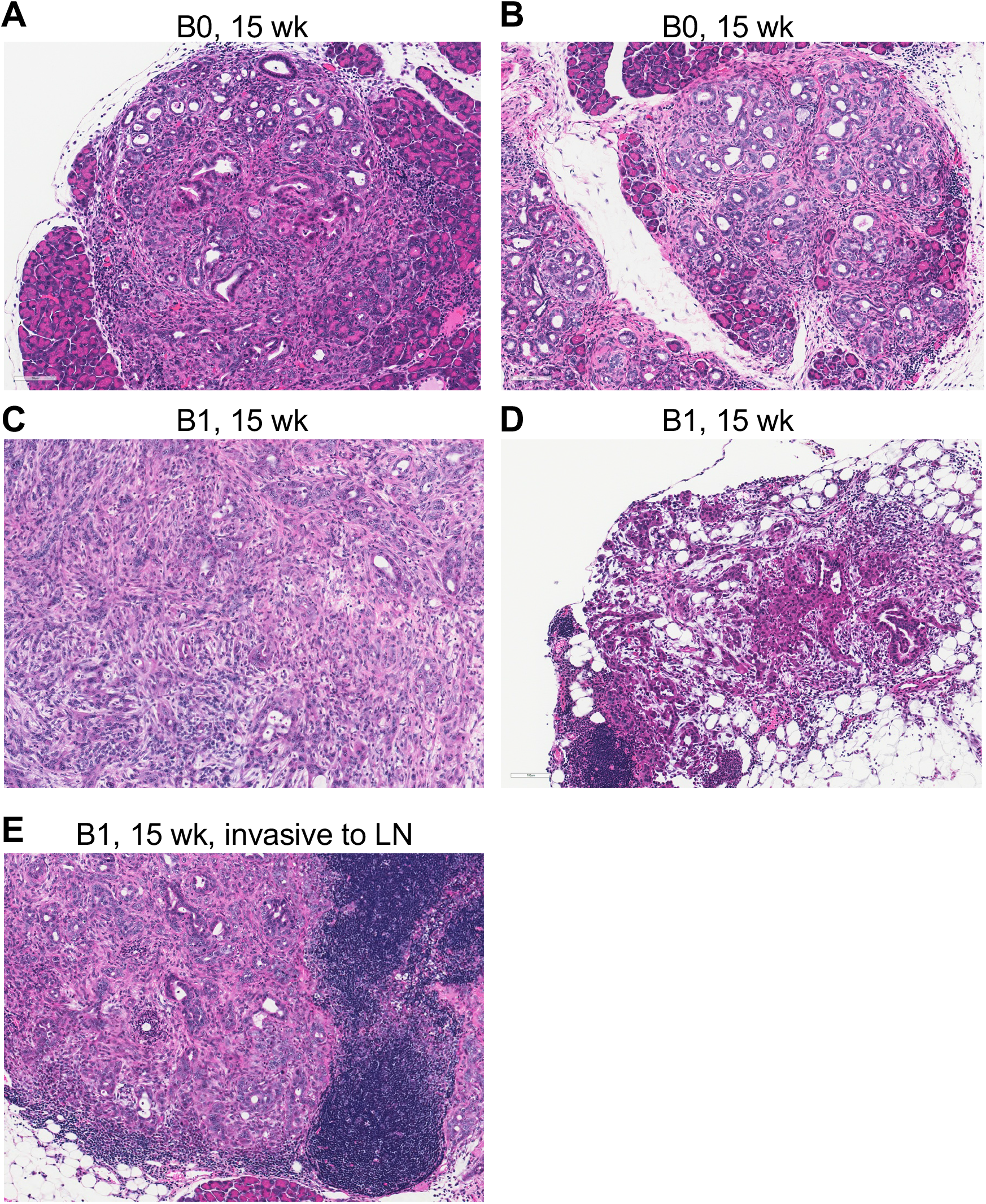
*HMMR*^*Δexon8-16*^ accelerated PDAC formation in *p48-Cre*; *LSL-KRAS*^*G12D*^; *p53*^*lox/+*^ mice. H&E stain of representative PDAC from B group mice at 15 weeks of age. (A and B) two B0: *p48-Cre*; *LSL-KRAS*^*G12D*^; *p53*^*lox/+*^ mice. (C and D) two B1: *p48-Cre*; *LSL-KRAS*^*G12D*^; *p53*^*lox/+*^; *HMMR*^*Δexon8-16/WT*^ mice, showing advanced PDAC (C) and early PDAC (D), respectively. (E) H&E stain of a peripancreatic lymph node (LN) invaded by PDAC from the same animal in (C). Original magnification, 200X.

### *p48-Cre*; *LSL-KRAS*^*G12D*^; *p53*^*lox/lox*^ mice had rapid PDAC progression irrespective of *HMMR*^*Δexon8-16*^ status

We established a cohort of group C mice of *p48-Cre*; *LSL-KRAS*^*G12D*^; *p53*^*lox/lox*^. These mice were further divided into 3 subgroups (C0, C1, and C2) based on the copy numbers of *HMMR*^*Δexon8-16*^. PDAC histology at the terminal stage among C0, C1, and C2 subgroups were similar. The C group mice developed advanced PDAC as the B group but at earlier ages (Figure 6A-C). The mice in the C group had very short life span (Figure 6D and Table 1, Log-rank test, C0: 67 mice, C1: 33 mice, and C2: 31 mice). The median survival ages for C0, C1, and C2 were all between 9 and 10 weeks (Table 1). There was no significant difference in survival found by log-rank test.

**Figure 6.**
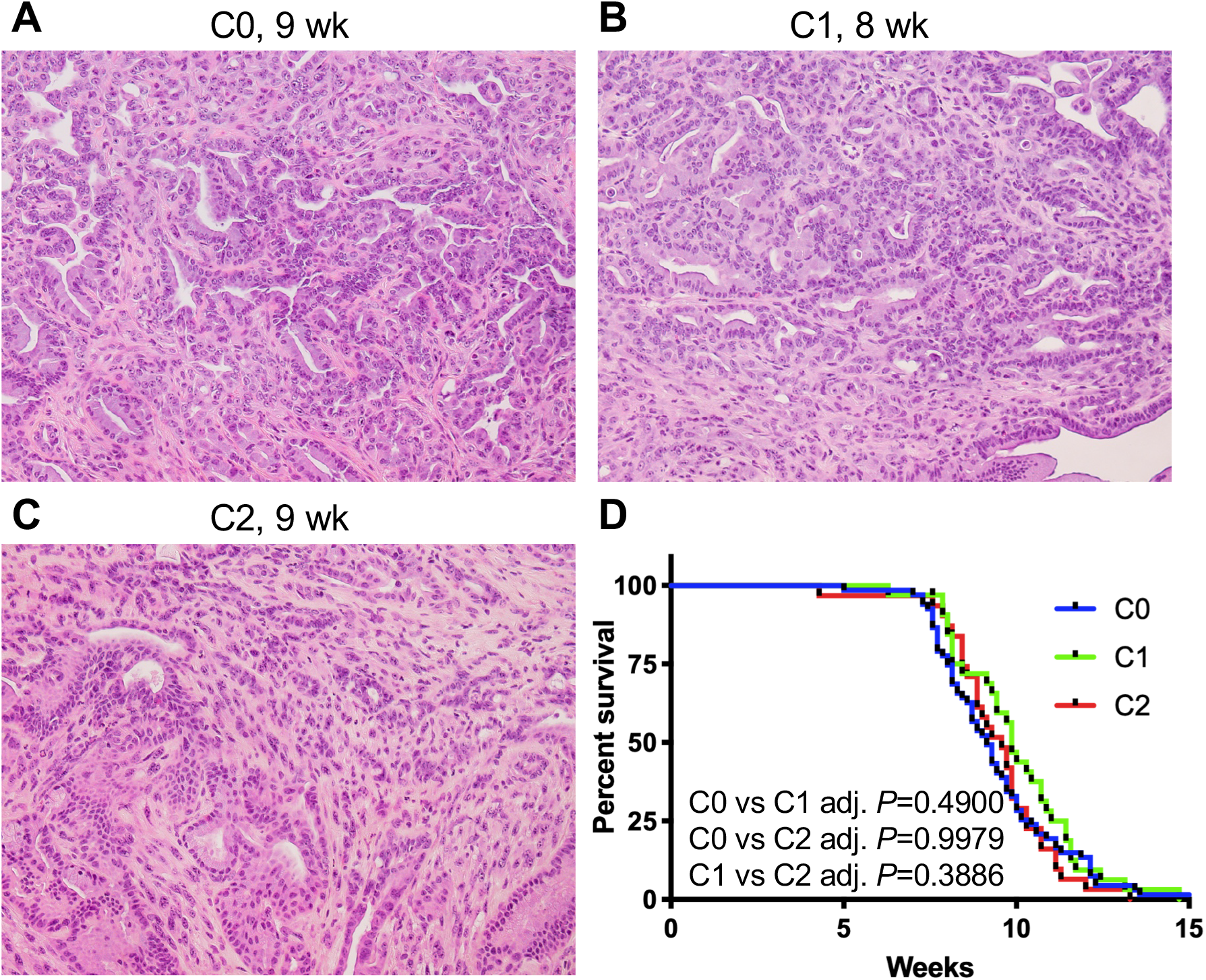
The *p48-Cre*; *LSL-KRAS*^*G12D*^; *p53*^*lox/lox*^ mice had too rapid PDAC progression to potentiate the effect of *HMMR*^*Δexon8-16*^. (A-C) H&E stain of representative PDAC from (A) a 9-week-old C0: *p48-Cre*; *LSL-KRAS*^*G12D*^; *p53*^*lox/ lox*^ mouse, (B) an 8-week-old C1: *p48-Cre*; *LSL-KRAS*^*G12D*^; *p53*^*lox/ lox*^; *HMMR*^*Δexon8-16/WT*^ mouse, and (C) a 9-week-old C2: *p48-Cre*; *LSL-KRAS*^*G12D*^; *p53*^*lox/ lox*^; *HMMR*^*Δexon8-16/Δexon8-16*^ mouse. (D) Kaplan-Meier survival curve for C0, C1, and C2 mice. Tukey adjusted *P* values from pairwise log-rank test were shown. Original magnification, 200X.

### *HMMR*^*Δexon8-16*^ reduced the survival of *RIP-Tag* PNET mice

*RIP-Tag* mice developed insulinoma, one subtype of PNETs [10]. As a result, the *RIP-Tag* mice died before 18 weeks of age due to hypoglycemia. Because homozygous *RIP-Tag* mice develop PNET faster [25], only hemizygous *RIP-Tag* mice in C57BL/6 background were used here. We established a cohort of *RIP-Tag* mice (T group) with different copy numbers of *HMMR*^*Δexon8-16*^ (T0, T1, and T2). Healthy mice have non-fasting blood glucose levels around 200 mg/dL, and T group mice developed PNET with blood glucose levels lower than 50 mg/dL at the end point (data not shown). PNET histology at the terminal stage among the T subgroups were similar (Figure 7A-C). *RIP-Tag*; *HMMR*^*Δexon8-16/WT*^ (T1) and *RIP-Tag*; *HMMR*^*Δexon8-16/Δexon8-16*^ (T2) died earlier than *RIP-Tag* (T0) (Figure 7D). Similar to the log-rank test results (Figure 7D), Cox regression analysis showed borderline significant difference between T1 vs T0: T1 is 46% more likely to die than T0 (*P* = 0.0573), and significant difference between T2 vs T0: T2 is 78% more likely to die than T0 (*P* = 0.0029). The median survival ages for T0, T1, and T2 mice were 17.57 weeks, 16.14 weeks, and 15.5 weeks, respectively (Table 1, T0: 58 mice, T1: 48 mice, and T2: 56 mice).

**Figure 7.**
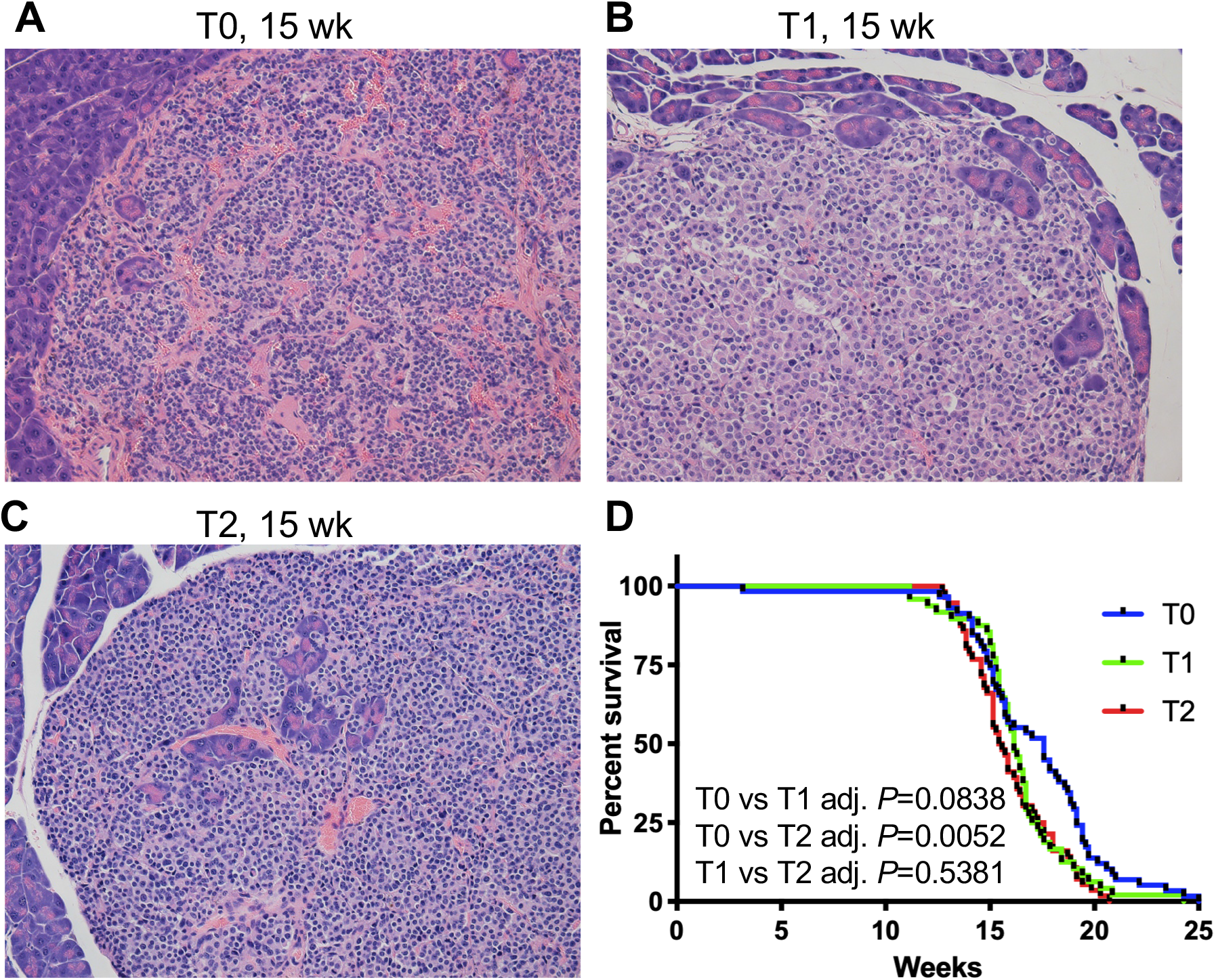
*HMMR*^*Δexon8-16*^ reduced the survival of *RIP-Tag* PNET mice. H&E stain of representative PNETs from (A) a 15-week-old T0: *RIP-Tag* mouse, (B) a 15-week-old T1: *RIP-Tag*; *HMMR*^*Δexon8-16/WT*^ mice, and (C) a 15-week-old T2: *RIP-Tag*; *HMMR*^*Δexon8-16/Δexon8-16*^ mouse. (D) Kaplan-Meier survival curve for T0, T1, and T2 mice. Tukey adjusted *P* values from pairwise log-rank test were shown. Original magnification, 200X.

### The association between *RHAMM* expression levels and *TP53* mutations or copy number loss in TCGA pancreatic cancer cohort

It was previously reported that TP53 transcriptionally downregulates *RHAMM* in colon carcinoma cell lines [18]. We evaluated whether the *RHAMM* expression is associated with *TP53* mutational status and copy number loss in pancreatic cancer patients using The Cancer Genome Atlas (TCGA) dataset that includes the data with gene expression (RNA-Seq), harmonized mutation, and gene level copy number. PDAC is the major pancreatic cancer type in the dataset. *TP53* mutations in this dataset include missense mutation, nonsense mutation, frame shift deletion, frame shift insertion, in-frame insertion, and splice site. We found upregulation of *RHAMM* in the pancreatic cancer patients with *TP53* mutations (Figure 8A, *P* < 0.05). Because *TP53* can also be inactivated by copy number loss, we evaluated whether the *RHAMM* expression is associated with the copy number variation (CNV) of *TP53*. We observed that *RHAMM* levels were higher in patients with one *TP53* copy number (CN=1) than in patients with 2 copy numbers (Figure 8B, *P* < 0.05). These data suggested that WT TP53 protein possess a transcriptional repression activity for *RHAMM/HMMR in vivo*. Then, we conducted a survival analysis using *RHAMM/HMMR* expression and *TP53* mutational status or its CNV. We found that patients have significant different survival outcome among different *TP53* status/*RHAMM* levels combination groups. Patients who have mutant *TP53*/high *RHAMM* had shortest survival among all four groups (Figure 8C, overall log-rank *P* = 0.024). Similarly, patients who have *TP53* CN=1/high *RHAMM* had shortest survival among four groups (Figure 8D, overall log-rank *P* = 0.026).

**Figure 8.**
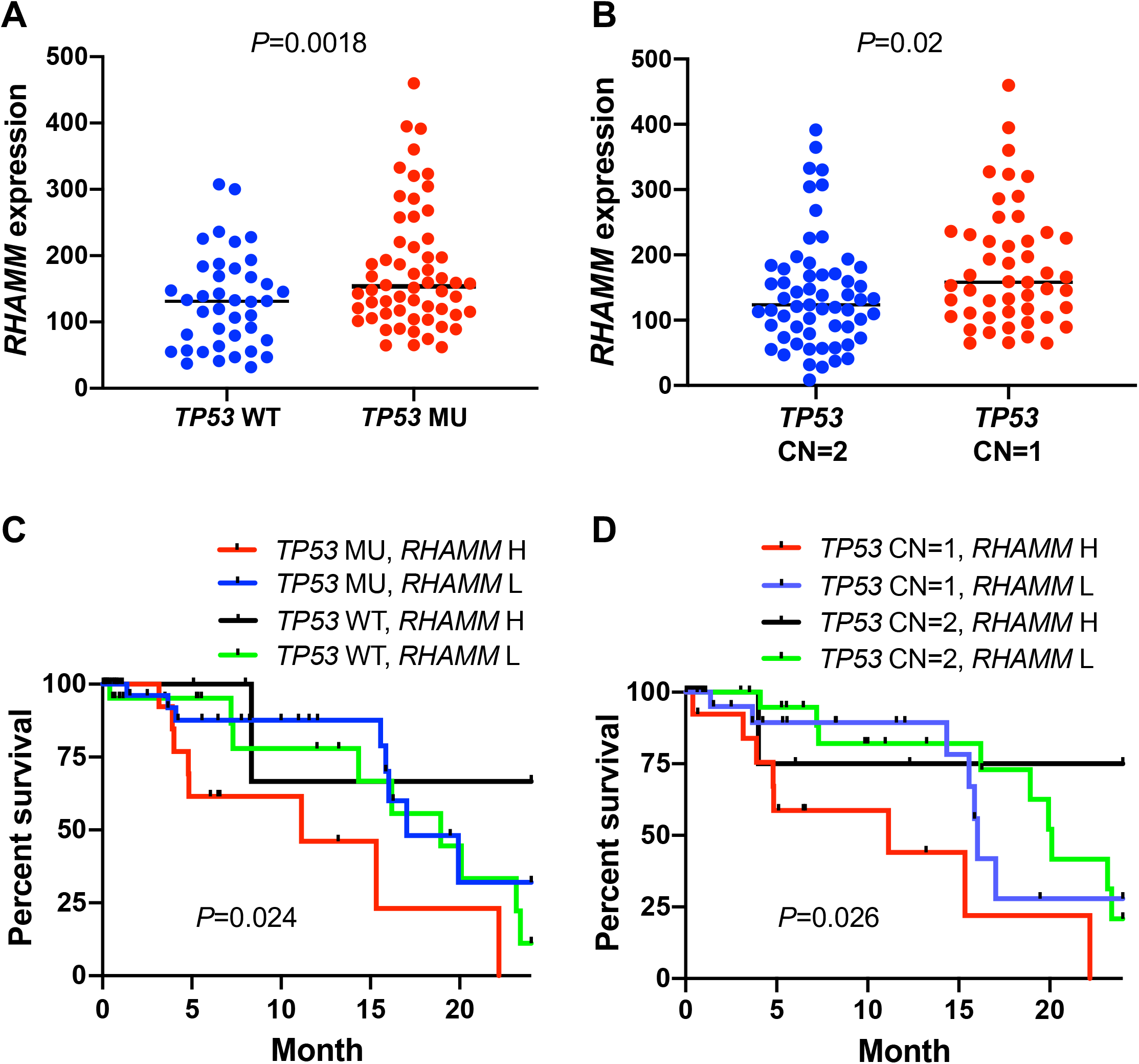
The association between *TP53* mutational status/copy number loss and *RHAMM/HMMR* expression levels in TCGA pancreatic cancer dataset. (A) Upregulation of *RHAMM/HMMR* expression in the TCGA pancreatic cancer patients with *TP53* mutations. WT: wild type (n = 53); MU: mutation (n = 66). (B) Upregulation of *RHAMM/HMMR* expression in the patients with copy number loss (CN =1, n =49) compared to autosomal copy number (CN =2, n = 59). (C) Combination of *TP53* mutations and high *RHAMM/HMMR* expression predicted worse outcome. (D) Combination of *TP53* copy number loss (CN =1) and high *RHAMM/HMMR* expression predicted worse outcome. High (H) and low (L) was based on the 75 quartile *RHAMM/HMMR* expression value. *P* values from log-rank test were shown in C and D.

## Discussion

Pancreatic cancer is lethal and new treatment options for this devastating disease are urgently needed. We have reported that upregulation of RHAMM/HMMR in human PDAC, PNET, and many other cancer types, while RHAMM /HMMR protein is not expressed in most of the normal tissues including pancreas [17, 20, 21, 26, 27]. This property of RHAMM makes RHAMM blockage a very attractive therapeutic target in pancreatic cancer.

To test the above idea in pre-clinical mouse models of PDAC and PNET, we originally took advantage of an existing RHAMM deletion strain, *Rhamm*^*-/-*^ mouse [23]. However, our molecular and histological characterization identified the expression of a truncated RHAMM (HMMR^Δexon8-16^) with a molecular weight of ∼27 kDa in the *Rhamm*^*-/-*^ mice. HMMR^Δexon8-16^ protein is more abundant than WT RHAMM/HMMR in WT mice. While HMMR^Δexon8-16^ by itself did not promote the progression of PanINs to PDAC (Group A, *p48-Cre*; *LSL-KRAS*^*G12D*^ mice), the combination of HMMR^Δexon8-16^ and heterozygous p53 loss promote earlier onset of invasive PDAC formation (Group B, *p48-Cre*; *LSL-KRAS*^*G12D*^; *p53*^*lox/+*^ mice). The more copies of *HMMR*^Δexon8-16^, the more likely that mice would die early for Group B mice and Group T, *RIP-Tag* PNET mice. Consistent with our findings using mouse models, pancreatic cancer patients who have mutant *TP53*/high *RHAMM* or *TP53* CN=1/high *RHAMM* had shortest survival among different *TP53* status/*RHAMM* levels combination groups. These data together supported that high levels of RHAMM/HMMR cooperated with dysfunctional p53 to accelerate the progression of pancreatic cancer in mice and in humans.

It has been shown that human RHAMM/HMMR is a G2/M protein [17, 18] and its C-terminus is recognized by the Anaphase-Promoting Complex (APC) for its degradation in the G1 phase [28]. Because this C-terminal homologous region is deleted in the mouse HMMR^Δexon8-16^ protein, the HMMR^Δexon8-16^ protein is likely not degradable by APC and this may contribute to the higher levels of the HMMR^Δexon8-16^ protein compared to the WT protein. Besides APC, two other E3 ubiquitin ligase activities, BRCA1/BARD1 and RFWD3, have been reported to regulate RHAMM/HMMR protein levels [29–31], but the consensus amino acid sequences for BRCA1/BARD1 and RFWD3 recognition are not known.

Exons 1 ∼ 4 of RHAMM/HMMR encode a microtubule-binding region, exons 5 ∼ 16 encode the central coiled-coil region, and exons 17 ∼ 18 encode a centrosome targeting region. Full-length RHAMM has 3 putative hyaluronan-binding sites (BX7B motif) (Figure 1A). HMMR^Δexon8-16^ protein retains one of the 3 putative hyaluronan-binding sites and exon 4, but it loses the C-terminal centrosome targeting region because the fusion of exon 7 and exon 17 resulted in a stop codon in the middle of exon 17. Our study indicated that the N-terminal microtubule-binding region and/or part of the central coiled-coil region are important for its oncogenic function, but the C-terminal centrosome targeting region is not. Further investigation is required to determine whether the remaining putative hyaluronan-binding motif is essential for the oncogenic function of HMMR^Δexon8-16^ and whether the lysine to threonine mutation at residue 71 affects any function. Between two of the alternative splicing human RHAMM isoforms, we have previously demonstrated that the RHAMM^B^, which does not have exon 4, is crucial for in vivo liver metastatic capacity of PNET, but RHAMM^A^, which carries exon 4, cannot promote liver metastasis of PNET [21]. It remains to be investigated whether a mouse RHAMM^B^ exists naturally or any mouse RHAMM isoform is upregulated in spontaneous mouse tumors. Because mouse HMMR^Δexon8-16^ protein still retains exon 4 and possesses an oncogenic function, we postulate that either loss of exon 4 or loss of the C-terminal centrosome targeting region confers the oncogenic function of RHAMM/HMMR.

In addition to *HMMR*^*Δexon8-16*^*/*^*Δexon8-16*^ (previously *Rhamm*^*-/-*^) mouse [23], *HMMR*^*m/m*^ and *HMMR*^*tm1a/tm1a*^ mice have been generated [32, 33]. A schematic of HMMR protein/gene in these mouse models can be found in the supplement figure 1 of [32]. The *HMMR*^*m/m*^ mice expresses HMMR^Δexon11-18^ protein with a molecular weight of 65 kDa and HMMR^Δexon11-18^ protein is also more abundant than WT proteins in mouse testis [33]. The homozygous *HMMR*^*m/m*^ mice are viable and normal in appearance [33]. However, HMMR^Δexon11-18^ causes hypofertility in older females (>24 weeks old) and in male mice independently of the male’s ages [31, 33]. Because *HMMR*^*m*^ (*HMMR*^*Δexon11-18*^) also loses C-terminal centrosome targeting region of RHAMM encoded by exons 17 ∼ 18 and some of central coiled-coil region encoded by exons 5 ∼ 16, *HMMR*^*m*^ (*HMMR*^*Δexon11-18*^) might promote invasive PDAC progression and reduce the survival of mice with PDAC and PNET similar to this study with *HMMR*^*Δexon8-16*^. A side-by-side comparison of *HMMR*^*Δexon11-18*^ and *HMMR*^*Δexon8-16*^*/*^*Δexon8-16*^ for their oncogenic property would help to further dissect the functional domains of HMMR. In contrast, *HMMR*^*tm1a/tm1a*^ mice only express the exon 1 and 2 of HMMR. *HMMR*^*tm1a/tm1a*^ mice produced litters with decreased survival, along with decreased adult body size and morphological defects in the brain, and very few mice survive to adulthood [32]. Thus, *HMMR*^*tm1a/tm1a*^ mice are not suitable for crossing to PDAC and PNET mouse models. A pancreas-specific knockout of *HMMR* would be needed to assess the effect of targeting RHAMM/HMMR in pancreatic cancer.

## Materials and Methods

### Mouse Strains, Animal Husbandry, and Genotyping

Cohorts of A, B, and C groups were of mixed genetic background. *p48-Cre; LSL-KRAS*^*G12D*^ mice were bred with *p53*^*lox/lox*^ mice and *HMMR*^*Δexon8-16/Δ exon8-16*^ mice (previously known as *Rhamm*^*-/-*^ [23]). Cohorts of PNETs are generated by crossing *RIP-Tag* mice (C57BL/6 genetic background) with and *HMMR*^*Δexon8-16/Δ exon8-16*^ mice. All procedures involving mice were approved by the Institutional Animal Care and Use Committee. There was no noticeable influence of sex on the results of this study. This study was carried out in strict accordance with the recommendations in the Guide for the Care and Use of Laboratory Animals of the National Institutes of Health. All mice were housed in accordance with institutional guidelines.

Mouse genomic DNA was prepared for PCR genotyping through incubation of each earpiece in 200 μl 0.05 M NaOH for 20 minutes at 98°C followed by the addition of 20 μl 1 M Tris-HCl (pH 7.5) at room temperature. Primers used in the PCR genotyping are as follows: for *HMMR*^*Δexon8-16*^: 5’-TGCTCGAGATGTCATGAAGG-3’ and 5’-CGAGAGGTCCTTTTCACCAG-3’ (a 360-bp product); for *HMMR* wild-type: 5’-CCAGTGCCCGAGAGAATTTA-3’ and 5’-TCCACTTGATCAGATGCACA-3’ (a 338-bp product) or 5’-AGGGAGCAGTACAGAGGTGT-3’ and 5’-TCGACAGCGTGTTCGGATAG-3’ (a 535-bp product); for *LSL-KRAS*^*G12D*^: 5’-CGCAGACTGTAGAGCAGCG-3’ and 5’-CCATGGCTTGAGTAAGTCTGC-3’ (a 650-bp product); for *p48-cre*: 5’-CTGATTTCGACCAGGTTCGT-3’ and 5’-ATTCTCCCACCGTCAGTACG-3’ (a 160-bp product); for *p53* allele: 5’-GGTTAAACCCAGCTTGACCA-3’ and 5’-GGAGGCAGAGACAGTTGGAG-3’ (a 370-bp product for *p53*^*lox*^, 240-bp product for WT). *RIP-Tag* PCR genotyping has been described [25].

### Sequencing of the *HMMR*^*Δexon8-16*^ Transcript and Western Blot Analysis

Mouse testes were removed and frozen on dry ice immediately after euthanasia. The testes were crushed into powder form with liquid nitrogen, and RNA was purified from the powder using the RNeasy Plus Mini Kit (QIAGEN, #74134). The purified RNA was used to synthesize cDNA using SuperScript^™^ IV VILO^™^ Master Mix with ezDNase^™^ Enzyme (Thermofisher, 11766050). An N-terminal HMMR fragment was amplified using AmpliTaq (Thermofisher, N8080153) or Q5 High-Fidelity DNA Polymerase (New England BioLabs, #M0491L) with the following primer pairs: (1) 5’-AACCAGAGCCAACGAGCTAC-3’ (on exon 5) and 5’-AGCCTTGGAAGGGTCAAAGT-3’ (on exon 17) (a 330 bp product to detect *HMMR*^*Δexon8-16*^); and (2) 5’-CTCCGGGTGCTTATGATGTT-3’ (on exon 2) and 5’-CCGTTTTTCCAGTGAAGCAT-3’ (on exon 5) (a 362-bp product for *HMMR*^*Δexon8-16*^ and WT to detect the missense mutation on exon 3). The resulting PCR products were purified using DNA Clean & Concentrator (Zymo, D4013) and were sent to PSOMAGEN, INC. (formerly Macrogen Corp.) along with the primer, 5’-CCGTTTTTCCAGTGAAGCAT-3’, for sequencing.

Frozen powders of mouse testes were lysed in NP-40 buffer (100 mM NaCl, 100 mM Tris pH 8.2, 0.5% NP-40) supplemented with a protease inhibitor mixture and PhosSTOP (Roche). Proteins were quantified by Bradford assay (Bio-Rad, Hercules, CA). Equal amounts of proteins were separated by SDS-PAGE and transferred to nitrocellulose membranes. To visualize equal protein loading, blots were stained with Ponceau S. Blots were incubated in 5% non-fat milk in TBST, probed with primary antibodies to RHAMM (clone EPR4054, Abcam, ab124729, 1:500) or Cyclin B2 (R&D, AF6204, 1 µg/mL), and then incubated with horseradish peroxidase (HRP)-conjugated secondary antibodies. Protein bands were visualized by enhanced chemical luminescence (Pierce, Rockford, IL).

### Immunohistochemistry

Mouse tissues were fixed in 10% buffered formalin overnight at room temperature and then transferred to 70% ethanol, paraffin-embedded, and sectioned. Slides were deparaffinized and rehydrated by passaging through a graded xylene/ethanol series before staining.

Immunohistochemistry was performed using rabbit anti-RHAMM antibody (clone EPR4054, Abcam, ab124729, 1:500) as the primary antibody, followed by color development using the VECTASTAIN Elite ABC kit and counterstaining with hematoxylin.

### Statistical Analysis

Mice from this study were monitored daily for signs of disease and mortality up to 1 year. Kaplan-Meier method and log-rank test was used to test the survival differences between groups. Simulation method was used to adjust for multiple comparisons for log-rank test.

Publicly available gene expression (RNA-Seq version 2), harmonized mutations and gene level copy number datasets from The Cancer Genome Atlas (TCGA) were downloaded from the Genomic Data Commons Data Portal (https://portal.gdc.cancer.gov). Mann–Whitney U test and survival analysis were conducted for this dataset.

All analyses were performed in statistical software GraphPad Prism (version 7.0e) or SAS Version 9.4 (SAS Institute, Cary, NC). *P* < 0.05 was considered statistically significant.

## Acknowledgements

We thank Cornelia Tolg and Eva Ann Turley for providing *Rhamm*^*-/-*^ mice. We thank Danny Huang, Annie Yang, Makheni Jean-Pierre, Lei Tan, Shuibing Chen, Bing He, Mai Ho, Leticia Dizon, Taotao Zhang, and Ruben Diaz for assistance. This work is partially supported by NIH R01CA204916-01A1, DoD W81XWH-16-1-0619, STARR I12-0043, and the Center for Translational Pathology at the Department of Pathology and Laboratory Medicine, Weill Cornell Medicine

## Abbreviations

APC: anaphase-promoting complex
CN: copy number
CNV: copy number variation
ES: embryonic stem
HMMR^Δexon8-16^: N-terminal RHAMM lacking exons 8 ∼ 16
MU: mutation
OS: overall survival
*p53*^*lox/+*^: heterozygous p53 loss
*p53*^*lox/lox*^: homozygous-null p53
PanIN: pancreatic intraepithelial neoplasia
PDAC: pancreatic ductal adenocarcinoma
PNET: pancreatic neuroendocrine tumor
PCR: polymerase chain reaction
RHAMM or HMMR: receptor for hyaluronic acid-mediated motility
RHAMM^B^: receptor for hyaluronan-mediated motility isoform B
Rb: retinoblastoma
RIP: rat insulin promoter
Tag: T antigen
TCGA: The Cancer Genome Atlas
WT: wild-type

